# The Phosphodiesterase NbdA Links c-di-GMP Signaling to Type IV Pili Function in *Pseudo-monas aeruginosa* PAO1

**DOI:** 10.64898/2026.03.20.713172

**Authors:** Anna Scherhag, Katrin Aras, Martina Ledermann, Jaqueline Rehner, Maike Karcher, Hanna Lang, Simone Stegmüller, Elke Richling, Nicole Frankenberg-Dinkel, Susanne Zehner

## Abstract

The phosphodiesterase (PDE) NbdA (NO-induced biofilm dispersion locus A) consists of a membrane-integrated MHYT domain, a degenerated diguanylate cyclase (DGC) AGDEF domain and an EAL domain. The integral membrane domain MHYT is proposed to sense a so far unknown extracellular signal and transfers the information to the cytosolic enzyme domains to modulate cellular c-di-GMP level. Here, we show that full length NbdA from *Pseudomonas aeruginosa* PAO1 is an active PDE *in vivo*. In line with its PDE activity, overexpression leads to slightly reduced global c-di-GMP levels, and reduced twitching motility.

Surprisingly, overexpression of truncated cytosolic NbdA variants exhibited increased c-diGMP levels, suggesting previously uncharacterized DGC activity despite lacking a canonical GGDEF motif. While full-length NbdA overexpression resulted in only slight c-di-GMP reduction, cytosolic variants induced a significant increase, indicating a potential for nonenzymatic effects like protein-protein interactions.

Further investigation revealed a connection between NbdA and type IV pilus (T4P) function. Overexpression of NbdA conferred resistance to the T4P-dependent phage DMS3vir, suggesting interference with T4P assembly or function. Microscopic analysis demonstrated dynamic localization of NbdA, partially co-localizing with T4P components, supporting a role in T4P regulation. However, no clear link was re-established with flagellar motor switching or chemotaxis signaling. These findings position NbdA in the complex signaling network of c-diGMP and T4P-mediated surface behavior in *P. aeruginosa*. Future work will focus on elucidating the precise mechanisms of NbdA’s PDE activity and its interplay with other DGC/PDE networks.

**Importance:** In this work, we show the *in vivo* activity of the membrane-bound phosphodiesterase NbdA of *Pseudomonas aeruginosa,* its role in c-di-GMP homeostasis, cellular localization and implications in surface behavior. Using strains overexpressing NbdA and truncated protein variants, we detected a strong defect in growth on solid surfaces and an altered phage susceptibility. Co-localization experiments supported further the hypothesis of interaction with the type IV pilus apparatus. We propose for NbdA to be part of the protein network responsible for c-di-GMP level modulation at the cell pole and thereby regulating the function of type IV pilus apparatus.

## Introduction

The bacterium *Pseudomonas aeruginosa* is ubiquitous in the environment such as in water and soil, and is capable of infecting plants, animals, and humans as an opportunistic pathogen (1). Especially in hospital environments, infections with *P. aeruginosa* have become a major health concern. The bacterium exhibits high intrinsic and mutational antibiotic resistance, which makes treatment of infections particularly challenging (2). Patients with weakened immune systems therefore often develop chronic infections, further exacerbated by the ability of *P. aeruginosa* forming biofilms.

The different lifestyles of *P. aeruginosa* as planktonic, free motile individual cells on one side, and the highly organized and specialized consortium of cells in a biofilm on the other side, allow the bacterium to thrive in fast-changing environments. The key molecule controlling the switch from planktonic cells to the biofilm state is the second messenger cyclic bis-(3′, 5′)dimeric guanosine monophosphate (c-di-GMP) (3). Many bacteria share the complex signaling network around c-di-GMP to control the lifestyle switch and many other cellular processes. The levels of the second messenger c-di-GMP inside bacterial cells are generally determined by two types of enzymes: diguanylate cyclases (DGCs) synthesizing the cyclic messenger molecule from two guanosine triphosphate (GTP) nucleotides and phosphodiesterases (PDEs) degrading it. Known DGCs possess a GGDEF domain, whereas PDEs possess either an EAL or a HD-GYP domain (3, 4).

*P. aeruginosa* encodes over 40 proteins with conserved domains presumably capable of synthesizing or degrading c-di-GMP (5–7). While most of them contain only one enzymatic domain - either a DGC or a PDE domain - sixteen proteins contain both. This redundancy of proteins in the cell suggests a very specialized function for the individual proteins. Most proteins possess signal receiver or transmission domains beside the enzymatic domains, that regulate the activity of the c-di-GMP-modulating enzymes further depending on environmental signals (6, 8). The concentration of c-di-GMP in the cell is further balanced through subcellular distribution and organization of these proteins in super modules, resulting in distinct local cdi-GMP pools (9–17). Many direct protein-protein interactions of DGCs and PDEs to other cdi-GMP modulating enzymes, effectors, cellular targets, sensoring, or regulating proteins have also been described (12, 18–21). Furthermore, so-called trigger PDEs were identified that modulate cellular functions independently of their enzymatic activity (9).

The membrane-integrated tandem phosphodiesterase NbdA (NO-induced biofilm dispersion locus A) is a multimodule protein, consisting of a membrane integrated MHYT-domain fused to a cytoplasmic DGC and PDE domain. The DGC domain possesses a degenerated AGDEF motif and the conserved EAL motif. *In vitro*-studies with heterologously produced and purified NbdA lacking the integral membrane MHYT domain demonstrated enzymatic PDE activity, but no detectable DGC activity (22). The role of NbdA in *P. aeruginosa* PAO1 remains unclear, while a PA14 transposon mutant of *nbdA* showed a clear phenotype in biofilm formation and motility (5). Although initial work from our group suggested the involvement of NbdA in nitric oxide (NO)-induced biofilm dispersion of PAO1 (22), more recent work has shown that a Δ*nbdA* strain was still capable dispersing biofilms upon NO exposure (23). Further reports suggested a function of NbdA in the regulation of the flagellar motor switch together with the PDEs DipA and RbdA (24). The PDE activity of these three enzymes was crucial for this effect since the output was mediated by the c-di-GMP effector protein MapZ. MapZ was shown to interact with the chemotaxis methyltransferase CheR1, thus regulating the methylation of chemoreceptors and flagellar motor switching (25). The three PDEs are thought to be located at the cell pole in the methyl-accepting chemotaxis protein (MCP) chemoreceptor array (24). A pull-down screening using NbdA as bait identified the type IV pili ATPase PilB as interaction partner (26). This protein stimulates the extension of type IV pili (T4P) (27, 28). Its interaction with NbdA indicates a potential link between this PDE with the type IV pili-mediated attachment or motility. Interestingly, in *P. aeruginosa* strain KT1115, NbdA was found to stimulate rhamnolipid synthesis during prolonged fermentation. Production of this biosurfactant was also directly linked to changes of c-di-GMP levels and the PDE activity of NbdA (29).

To further understand the role of NbdA in cellular function, we used a gain-of-function approach by overexpressing *nbdA* in *P. aeruginosa* PAO1. For this purpose, we created PAO1 strains that overproduce all subdomains of the protein individually and in different combinations, as well as enzymatic inactive variants to test for putative moonlighting effects.

## Material and Methods

### Cell cultivation

All strains and plasmids used in this study are listed in Table S1 and S2. Cells were grown in baffled flasks in LB medium (Carl Roth, Karlsruhe, Germany) for 16–18 h at 37°C, 180 rpm. The medium was supplemented with antibiotics when required for *E. coli*: 100 µg/mL ampicillin, 10 µg/mL gentamicin and 5 µg/mL tetracycline; for *P. aeruginosa* 300 µg/mL gentamicin and 150 µg/mL tetracycline.

### Generation of markerless mutations, overexpression and site-specific mutagenesis

Markerless deletion mutants in *P. aeruginosa* PAO1 were generated by two step allelic exchange according to (30). The DNA fragments for gene deletion were created by splicing by overlap extension PCR and integrated into the vector pEXG2 by Gibson assembly. The deletion plasmids were introduced into *P. aeruginosa* PAO1 by biparental mating (31). Exconjugants were selected on antibiotic gentamicin. Subsequently, exconjugants were plated on sucrose-containing media for counterselection against *sacB* to force the excision of the vector backbone from the integration site.

For overexpression of *nbdA* and truncated gene variants in *P. aeruginosa* PAO1 specific PCR fragments were generated and ligated in pHerd26T by Gibson assembly. QuikChange^TM^ mutagenesis was performed to introduce specific amino acid exchanges in the sequence of NbdA. All primers used in this study are listed in Table S3.

### *In vivo E. coli* complementation assay

PCR fragments for *nbdA* and truncated gene fragments were cloned in pCYB1 vector using Gibson cloning (Table S2 and S3) and introduced into *E. coli* AB607 (Table S1). Overnight cultures in LB with antibiotic selection were diluted to an OD_600_ = 1 in swimming medium (5 g/L NaCl, 10 g/L trypton). 1 µL of each strain was dropped on swimming agar plates (0.3% (w/v) Difco-Agar (BD), 5 g/L NaCl, 10 g/L tryptone) supplemented with 100 µM IPTG. Plates were incubated for 5 h at 37°C.

### Overproduction of protein variants in *P. aeruginosa*

Phenotypic assays were carried out with *P. aeruginosa* strains containing expression vectors for protein variant overproduction (Table S2). Precultures of desired strains were prepared in LB medium supplemented with tetracycline and were diluted 1:20 in fresh medium. Cultures were grown at 37°C shaking to an OD_600_ = 0.5. Gene expression was induced by the addition of 0.1% (w/v) L-arabinose and cultures were incubated at 22°C shaking for 5 h.

### Chemicals and standards for quantification of nucleotides

The standards used for the calibration curves, GMP (>99%), c-di-GMP (≥95%) and pGpG (≥95%), were purchased from Sigma-Aldrich (St. Louis, USA), Biomol (Hamburg, Germany) and Jena Bioscience (Jena, Germany), respectively. Isotopically labelled GMP (^15^*N* ^13^*C* -GMP, >95%) was obtained by Silantes (München, Germany) and ^15^*N* ^13^*C* -c-di-GMP (≥95%) by BioLog (Bremen, Germany). MS grade ammonium acetate (>99%) and MS grade methanol were purchased from Sigma-Aldrich (St. Louis, USA) and acetic acid (LC-MS grade) from Bio-solve (Valkenswaard, Netherlands), respectively.

### Sample preparation and mass spectrometric quantification of GMP, c-di-GMP, and pGpG in lyophilized bacteria extracts

Extraction and quantification of nucleotide messengers was modified after (32). For nucleotide extraction, *P. aeruginosa* cultures overproducing the protein variants (Table S1) were used. The OD_600_ was adjusted to 1.5 in 5 mL volume. Cell pellets were washed twice in LB and subsequently resuspended in 300 µL extraction solvent (acetonitrile:methanol:water 2:2:1). The samples were incubated on ice for 15 min, followed by 10 min at 95°C and centrifuged for 10 min at 4°C at 20.800 x g. The supernatant was transferred to a new precooled test tube. A second and third extraction was carried out by adding 200 µL extraction solvent to the pellet, resuspension, and incubation on ice for 15 min. After each extraction, tubes were centrifuged at 20.800 x g for 10 min and the supernatant was collected. The extracts (700 µL each) were incubated at−20°C for 16 h to precipitate leftover protein, centrifuged again, and the supernatant was lyophilized at −80°C and at 0.04 mbar (freeze and dry machine LSC plus, Christ, Osterode, Germany). Immediately before the analysis, the lyophilized bacteria extracts were resuspended in water (200 µL) and centrifuged (17,000 x g, 10 min, 4°C). 40 µL of the supernatant were transferred in 1.5 mL vials with inserts and 40 µL of internal standard mix (4 µM ^15^*N*_5_^13^*C*_10_-GMP and 10 nM ^15^*N*_10_^13^*C*_20_-c-di-GMP) were added, so that the final concentrations of the labelled standards in the vial were 2 µM for ^15^*N*_5_^13^*C*_10_-GMP and 5 nM for ^15^*N*_10_^13^*C*_20_-c-di-GMP. The samples were mixed and used for analysis in duplicates, which was carried out on an Agilent 1290 HPLC, consisting of a binary pump (G4220A), a column oven (TCC G1316C), an autosampler (G4226A), and a thermostat (G1330B) (Agilent Technologies, Waldkirch, Germany), coupled with a QTrap 5500 tandem mass spectrometer (AB Sciex, Darmstadt, Germany). Separations were performed using a reversed phase column (Luna C8, 150×3 mm, 3 µm, Phenomenex, Aschaffenburg, Germany) equipped with a respective guard column at 30°C. The mobile phases consisted of 50 mM ammonium acetate at pH 4.57 (A) and methanol (B) at a flow rate of 400 µL/min. The gradient program was as follows: 1% B at 0 min, held for 2 min, then raised to 3% over 0.1 min and further raised to 20% till 6 min. Next, the content of B was set to initial conditions over 0.5 min with a following reconditioning step. 20 µL injection volume were used for the analysis of c-di-GMP and pGpG, and 0.1 µL for the analysis of GMP, due to higher amounts of GMP in the samples. The measurement was carried out in the scheduled multiple reaction monitoring (sMRM) mode with a detection window of 100 sec. For ionization an electrospray source (ESI), operated in positive mode, was used. The source parameters were as follows: ion spray voltage 1500 V, temperature 550°C, nebulizer gas 35 psi, heater gas 55 psi, curtain gas 20 psi. The compound specific parameters are listed in Table S4. For each analyte and the two isotopically marked standards, two mass transitions (Q1→Q3, *m*/*z*) were measured, first (marked with an asterisk) for quantification and second as confirmatory signal: GMP 363.9→151.9*, 363.9→134.9; c-di-GMP 690.9→152.1*, 690.9→539.9; pGpG 709.1→152.0*, 709.1→558.1; ^15^*N*_5_^13^*C*_10_-GMP 378.9→162.0*, 378.9→144.0; ^15^*N*_10_^13^*C*_20_-c-di-GMP 720.9→162.0*, 720.9→560.1. The limit of detection (LOD) and limit of quantification (LOQ) for the described method are 10 nM and 20 nM for GMP and 0.01 nM and 0.1 nM for c-di-GMP and pGpG, respectively.

For quantification of GMP a calibration curve with peak area ratio of GMP/^15^*N*_5_^13^*C*_10_-GMP against the concentration ratio of GMP/^15^N_5_^13^C_10_-GMP was created. For the quantification of both, c-di-GMP and pGpG ^15^*N*_10_^13^*C*_20_-c-di-GMP was used as internal standard. The calibration curves were in a range of 1–25 µM for GMP with a concentration of 2 µM ^15^*N*_5_^13^*C*_10_-GMP. The range for c-di-GMP and pGpG was 0.5–1000 nM and 0.5–25 nM, respectively, with a concentration of 5 nM of the internal standard ^15^*N*_10_^13^*C*_20_-c-di-GMP. MS data were evaluated by Analyst version 1.7.2 (AB Sciex, Darmstadt, Germany). From the linear regressions, the amounts of GMP, c-di-GMP and pGpG were deduced. The quantitative contents of GMP, c-di-GMP and pGpG in the bacterial extracts were calculated for three biological replicates referring to the protein content.

### Adhesion assay

*P. aeruginosa* was cultured in BM2 medium (7 mM (NH_4_)_2_SO_4_, 40 mM K_2_HPO_4_, 22 mM KH_2_PO_4_, 2 mM MgSO_4_, 0.4% (w/v) glucose, 0.01 mM FeSO_4_, pH 7) over night at 37°C 120 rpm. The cell density was adjusted to an OD_600_ = 1.5 in adhesion medium (BM2 medium containing 0.5% (w/v) casamino acids) containing tetracycline and 0.1% (w/v) L-arabinose. A round bottom 96-well plate was inoculated with 100 μL culture per well. The 96-well plate was incubated in a moist chamber for 1 h at 37°C. After incubation, the liquid cultures were discarded, and each well was washed with 300 μL distilled H_2_O. The attached biomass was stained with 125 μL of crystal violet solution (0.1%) for 15 min. Wells were then washed twice with 150 μL H_2_O and dried for 5 min at room temperature. The bound crystal violet was resuspended by the addition of 200 μL 95% (v/v) ethanol to each well and incubated for 15 min. The absorption at 595 nm was measured in a TECAN plate reader and the relative adhered biomass was estimated.

### Twitching assay

Precultures of *P. aeruginosa* strains were prepared in 8 mL LB medium supplemented with antibiotic and grown overnight at 37°C and 160 rpm. Cultures were then diluted 1:20 in 8 mL of LB medium and further incubated at 37°C and 160 rpm up to OD_600_ ∼ 0.5. Gene expression was induced by the addition of 0.1% L-arabinose and further incubated. After 5 h, cell density was adjusted to an OD_600_ = 1.5. Twitching-agar was supplemented with an antibiotic (100 µg/mL tetracycline) and inductor (0.1% -L-arabinose). The agar was then pierced with a sterile Pasteur pipette and 1 μL culture was spotted on the bottom of the petri dish. Plates were incubated for 40 h at 37°C. Subsequently, the agar medium was carefully removed, and the twitching zones at the bottom of the petri dish were stained with 0.05% (w/v) crystal violet for 20 min. The staining solution was carefully poured off; the petri dishes were rinsed with sterile distilled water and dried at room temperature. Images of the plates were taken and the diameter of the twitching zone was determined using ImageJ/GIMP software.

### Confocal fluorescence microscopy

The cellular localization of NbdA was investigated in *P. aeruginosa* cells containing genomically integrated and plasmid encoded fusions of NbdA to the fluorescent protein mNeon-Green. For protein detection, a C-terminal His_6_ tag was attached to the fusions. Cultures of strains with a genomically encoded *nbdA-*mNeonGreen fusion were prepared in 20 mL LB medium and grown at 37°C for 16 h. The culture was diluted 1:4 in fresh LB medium. Cultures of strains containing pME6032-*nbdA*-mNeonGreen were diluted 1:100 in 10 mL fresh LB with antibiotic and grown at 37°C shaking. When cultures reached an OD_600_ of 0.2–0.5, target gene expression was induced with 100 µM IPTG. Cultures were further incubated for 2 h at 37°C. For CLSM 3 μL culture were spotted on an agarose pad (1.2% w/v agarose in PBS, 1 mm) air dried for 5 min and covered. Fluorescent NbdA fusions were excited at 488 nm and fluorescence detected from 500–550 nm.

### Co-localization of NbdA with flagella, chemotaxis proteins and pili

Co-localization assays of NbdA-mNeonGreen with the polar flagellum of *P. aeruginosa*, the flagellar subunit FliC was modified. The introduction of a cysteine residue into FliC at the position 394 allows flagellar staining with the red fluorescent dye Alexa Fluor™ 594 C5 Maleimide (Thermo Fisher Scientific, Darmstadt, Germany). The staining protocol of Zhu *et al.* was modified as follows (33). Cultures were grown overnight at 37°C and 160 rpm in flasks containing 10 mL LB medium. Cells of an equivalent of OD_600_ = 0.5 in 1 mL were spun down at 1,000 x g for 5 min. The cell pellet was carefully resuspended in 50 μL PBS buffer containing 50 μg/mL Alexa Fluor™ dye and incubated for 15 min in the dark. Cells were collected by a 5 min centrifugation step at 1,000 x g and washed carefully in 300 μL 1x PBS buffer. Cells were centrifuged again for 5 min at 1,000 x g and then resuspended in 30 μL 1x PBS buffer for microscopy. 3 μL of stained cells were pipetted on an agarose pad air dried and covered. The histidine kinase component CheA of the chemotaxis machinery of *P. aeruginosa* locates to the flagellated cell pole and is present in nearly all cells (34). Therefore, CheA was used in co-localization studies with NbdA to mark the flagellated pole. Fusions of *cheA* to *tdtomato*, encoding for a red fluorescent protein was genomically integrated into the original gene locus. The *nbdA-mNeonGreen* fusion was plasmid-encoded and under the control of an inducible *tac* promoter.

Cultures of strains containing pME6032-*nbdA*-mNeonGreen were diluted 1:100 in 10 mL fresh LB containing antibiotic and grown at 37°C shaking until the cultures reached an OD_600_ of 0.2–0.5. Target gene expression was induced with 100 µM IPTG. Cultures were further incubated for 2 h at 37°C for the co-localization with flagella and CheA, and 5 h at 22°C for co-localization with PilO. For CLSM 3 μL of the respective cultures were spotted on a cover slip and covered with an agarose pad (1.2% w/v agarose in PBS, 1 mm). Fluorescent proteins were detected using excitation for Alexa Fluor^TM^: at 594 nm, mCherry at 594 nm, mNeon-Green: 488 nm, tdTomato: 514 nm. The fluorescence emission of each fluorescent protein was detected in a separate track.

### Phage infection assay

*P. aeruginosa* is infected by bacteriophage DSM3*vir*. The infection by DSM3*vir* is dependent on the exposure of type IV pili at the cell surface (35). Phage infection assays were performed to check whether *P. aeruginosa* type IV pili production is affected by the overexpression of *nbdA*. Therefore, bacteria were grown in LB supplemented with tetracycline at 37°C. A preculture was diluted in LB to an OD_600_ of 0.05. A flat-bottom 96-well plate was inoculated with 200 µL bacterial suspension in each well and incubated in a TECAN plate reader at 37°C shaking. The OD_595_ was measured every 30 min for 24 h. Gene expression was induced by addition of 0.1% (w/v) L-arabinose. One hour after induction, 10^5^ PFU/mL were added. Phage stocks and phage titer were determined as previously described (36).

### Statistics

Experimental data was tested for significant changes. Only biological replicates were considered for statistic evaluation. Technical replicates were averaged. If not stated otherwise, average and standard deviation of biological replicates were calculated with the respective formula in Excel (version 2112, Microsoft, Redmond, USA). To determine which statistical test is appropriate, data was tested for normal distribution in Origin according to Shapiro-Wilk or Kolmogorov-Smirnov. When data was normally distributed, a Student’s *t*-test was performed in Excel. The tail was dependent on the tested hypothesis and if the datasets were compared for changes (two-tailed test) or for higher/lower values (one tailed test). The type of *t*-test was determined with *F*-test as a test for equal variances, which was also performed in Excel. If the result of the *F*-test was > 0.05, data was homoscedastic (type 2), if the *F*-test < 0.05, the data was heteroscedastic (type 3). For paired datasets, type 1 was used. The significance level was 0.05, with 0.01 < *p* < 0.05 (*), 0.001 < *p* < 0.01 (**) and *p* < 0.001 (***). For multiple comparisons, a *p*-value correction according to Bonferroni was carried out.

## Results

### NbdA is an active phosphodiesterase in a heterologous host

Although it has already been shown that the cytoplasmic domains of NbdA (NbdA-AGDEF-EAL) possess PDE activity, experimental proof for the full-length protein is still lacking (22). We therefore used a functional complementation assay to test the activity of the full-length protein *in vivo*. The *E. coli* mutant (Δ*pdeH*) is deficient in swimming motility due to the deletion of the master phosphodiesterase PdeH (37). Complementation with an active PDE should restore swimming motility (Fig. 1). Plasmids encoding NbdA and truncated variants were transformed into this *E. coli* mutant and tested for swimming motility. Complementation with the full-length NbdA (MHYT-AGDEF-EAL) and the cytosolic NbdA (AGDEF-EAL) variant was able to rescue the *E. coli* Δ*pdeH* swimming deficiency similarly to the homologous complementation with the *E. coli* protein PdeH (Fig. 1). A single amino acid exchange in NbdA at the conserved EAL motif (MHYT-AGDEF-**<u>A</u>**AL) rendered the protein inactive. This variant was unable to restore the swimming activity of the *E. coli* Δ*pdeH* mutant strain. We therefore confirmed that full-length NbdA (MHYT-AGDEF-EAL) is an active PDE in a heterologous host. Interestingly, truncated variants of NbdA containing only the PDE domain (*nbdA*-EAL) failed to complement *E. coli* Δ*pdeH* suggesting a crucial role of the MHYT and AGDEF domains for the enzymatic activity.

**FIG. 1.**
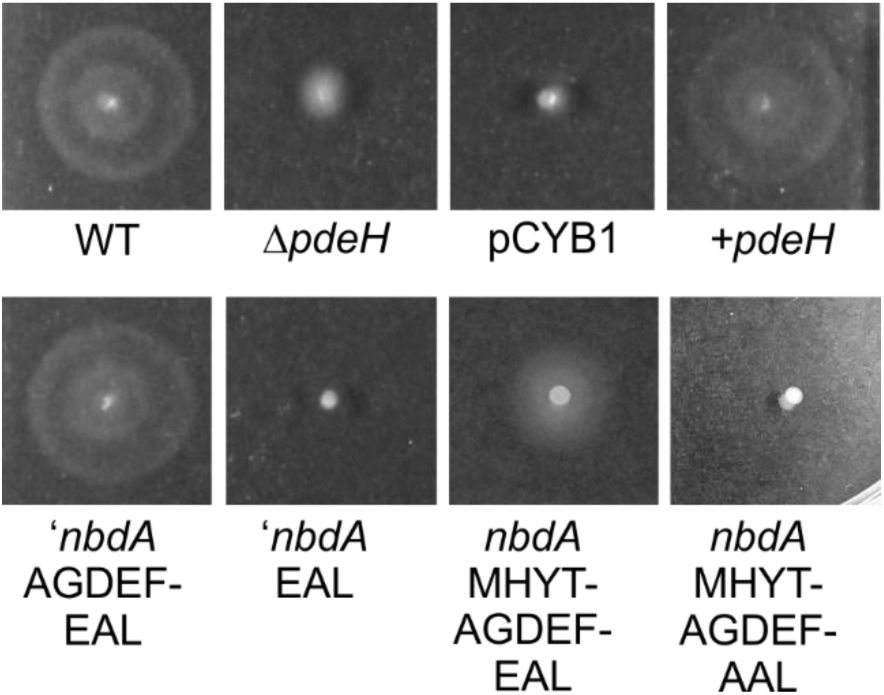
*Escherichia coli* Δ*pdeH* complementation assay. Swimming motility of parental *E. coli* MG1655 (WT), deletion strain *E. coli* AB607 (Δ*pdeH*), and complemented Δ*pdeH* strains with pMRE01 (+*pdeH*), empty vector (pCYB1), pMRE04 (*nbdA*_MHYT-AGDEF-EAL), pMRE09 (*nbdA*-MHYT-AGDEF-AAL), pMRE02 (*nbdA*-AGDEF-EAL), pMRE03 (*nbdA*-EAL) is shown. Cultures were inoculated on soft agar plates containing 100 µM IPTG and incubated at 37°C in a moist chamber for 5 hours. All experiments were tested in three biological replicates with 5 technical replicates.

### Expression of *nbdA* and variants in *Pseudomonas aeruginosa* PAO1 affects the second messenger c-di-GMP concentration

While the functional complementation confirmed a PDE activity for NbdA, phenotypic analyses of a PAO1 Δ*nbdA* mutant did not reveal any obvious phenotype regarding swimming motility or biofilm formation compared to the wild type (Fig. S1). Therefore, a gain-of-function approach was chosen. Plasmids for inducible expression of *nbdA* variants were introduced into *P. aeruginosa* PAO1 Δ*nbdA*. The full-length protein and the truncated NbdA variants for single and tandem domains were produced with a C-terminal *Strep* II-tag in PAO1 Δ*nbdA* (Fig. S2). All strains showed similar growth behavior in liquid medium, independent of the presence or absence of the inducer (Fig. S3).

To test whether NbdA shows PDE activity in these overexpression strains, we assessed cellular c-di-GMP levels and total nucleotide second messenger content. The nucleotides c-di-GMP, pGpG, and GMP were quantified by stable isotope dilution analysis (SIDA). For strains overexpressing *nbdA* variants, we observed two patterns in global concentrations of c-di-GMP. The first pattern, which was observed for MHYT domain-containing variants, showed low levels of c-di-GMP (6–23 pmol/mg protein) similar to the wild-type strain. The second pattern was the opposite; expression of the cytosolic variant lacking the MHYT domain showed a 12 to 40-fold increase in c-di-GMP content (>280 pmol/mg protein), (Fig. 2A). This result was intriguing, as the cytosolic variant of NbdA was previously shown to possess phosphodiesterase activity *in vitro* and in a heterologous host. Based on these findings, reduced cellular c-di-GMP levels had been anticipated. We further analyzed the cellular c-di-GMP level for strains producing enzymatically inactive NbdA variants. A full-length variant, where functional residues in the PDE domain were exchanged (NbdA-MHYT-AGDEF-<u>AAL</u>), increased cellular c-di-GMP levels (>2.5 fold) compared to the control strain (PAO1 with empty vector) and the strain overexpressing the native *nbdA* sequence. This result is consistent with the suggested function for the modified/inactive PDE domain. For the cytosolic domains of NbdA, two inactive protein variants were produced, the DGC ^−^ variant “AGAAF-EAL” and the PDE ^−^ variant “AGDEF-AAL”. The overexpression of these inactive cytosolic domains did not alter the cellular c-di-GMP levels compared to the unmodified cytosolic variant “AGDEFEAL”.

**FIG. 2.**
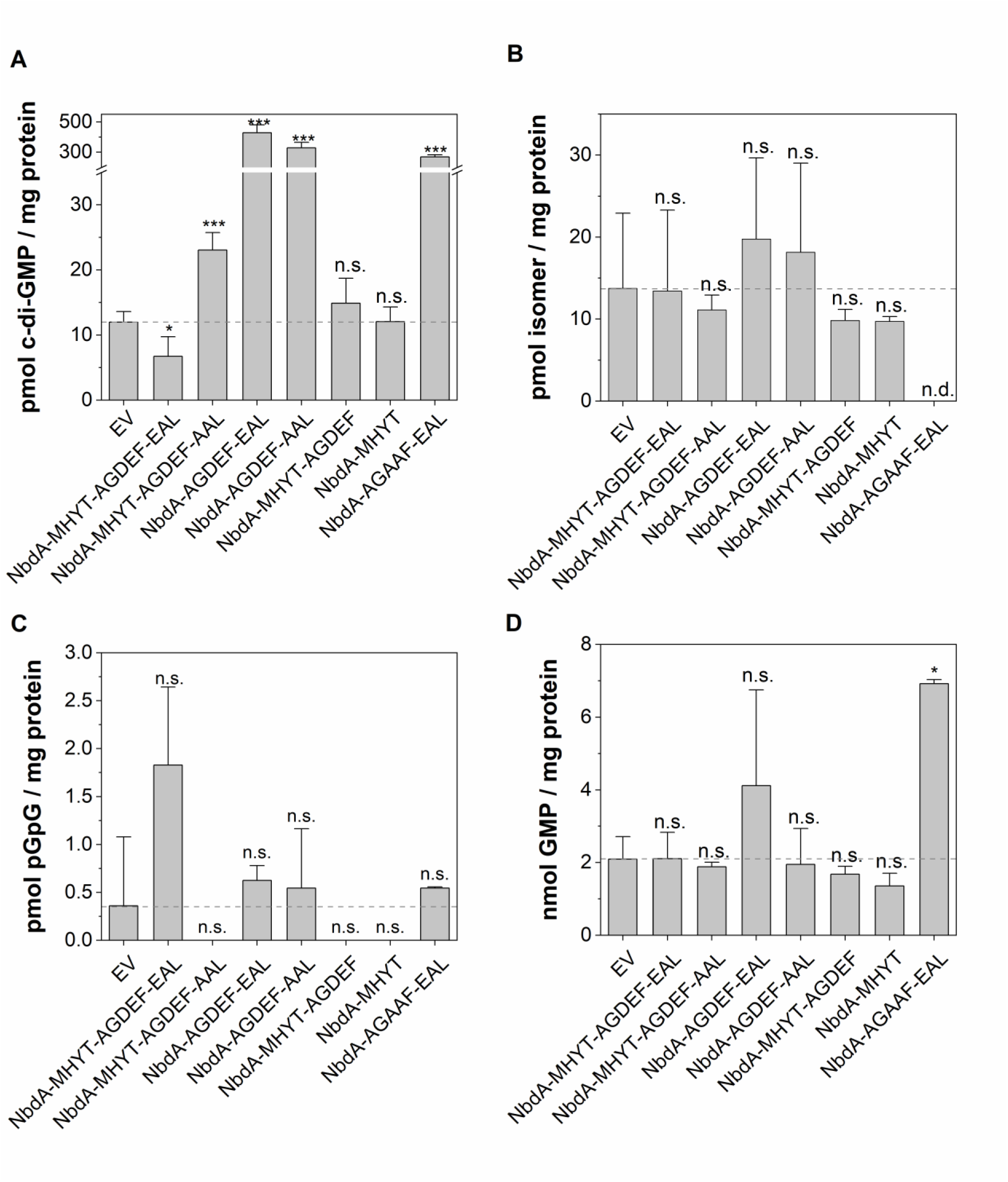
Quantification of nucleotide second messengers in cell extracts from *P. aeruginosa ΔnbdA* strains overproducing NbdA variants by HPLC-ESI_pos_-MS/MS using stable isotope dilution analysis (SIDA). The concentration of c-di-GMP (A), isomer of c-di-GMP (B), pGpG (C) and GMP (D) in the extracts are shown as mean ± standard deviation (SD) for three biological replicates, referring to the total protein content. Significant changes are marked with asterisks, determined by the student’s t-test. *** p<0.001, * p<0.05, n.s. not significant, n.d. not determined.

The degradation products of c-di-GMP, the nucleotides pGpG and GMP, were detected over-all in very low concentrations in the cell extracts (Fig. 2C, D). The determined concentrations did not differ significantly between overexpressing strains, wildtype, and vector control.

Interestingly, a so far unidentified c-di-GMP isomer was detected in all tested cell extracts of overexpression strains (Fig. 2B), but also in in the cell extracts of PAO1 and the *nbdA* deletion strain (Fig. S1). The isomer revealed the same mass to charge (*m/z*) ratio as c-di-GMP but showed a different fractionation pattern. The compound was tested for being the frequently occurring (2’5’)-(3’5’)-c-di-GMP isomer, short 2’3’c-di-GMP. Therefore, standards were ordered and analyzed. The unknown c-di-GMP isomer from the cell extracts did not resemble properties with the 2’3’-c-di-GMP standard. The measurements of the compound for all tested strains showed high variations and was likely independent of NbdA variant overproduction. Therefore, we didn’t pursue the identification of the structure of this compound.

### Overexpression of *nbdA* leads to reduced growth on solid surfaces

Consistent with the previous observations showing normal growth in liquid medium, the PAO1 Δ*nbdA* strain harboring the plasmid for inducible expression of native *nbdA* displayed normal growth on solid medium in the absence of the inducer L-arabinose. In contrast, induction of *nbdA* expression resulted in reduced growth on solid medium (Fig. 3).

**FIG. 3.**
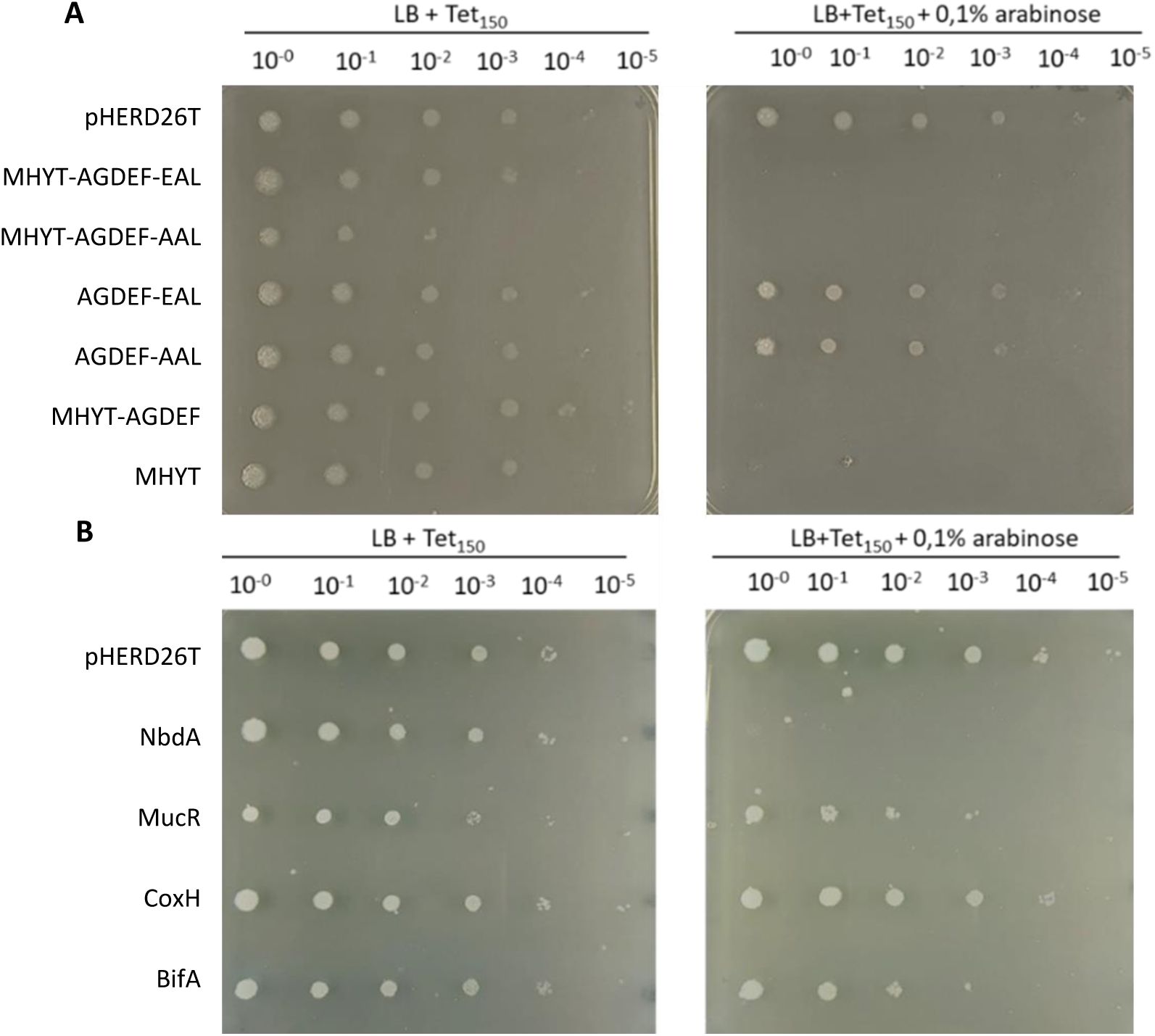
Growth on solid media tested by drop dilution assay. **A**) *P. aeruginosa* PAO1 Δ*nbdA* containing pHerd26T derivatives for overproduction of NbdA protein variants were grown in a liquid preculture. The OD was adjusted to 0.1 and dilution series were prepared in fresh medium. From this preculture 1 µl of each dilution was spotted on 1.5% LB agar with and without inductor L-arabinose. PAO1 Δ*nbdA* with empty vector (pHERD26T), pMRP12 (MHYT-AGDEF-EAL), pMRP14 (MHYT-AGDEF-AAL), pASC04 (AGDEF-EAL), pMKE02 (AGDEF-AAL), pJRE03 (MHYT-AGDEF), pJRE02 (MHYT). **B)** *P. aeruginosa* PAO1 Δ*nbdA* was transformed with plasmids encoding the membrane proteins NbdA, MucR, and BifA of *P. aeruginosa* PAO1, and CoxH of *Afipia carboxidovorans* under the control of pBAD promoter. The control strain harbours the empty vector (pHERD26T). From precultures without inductor, the cells were diluted and subsequently inoculated on 1.5% LB agar with and without the inductor L-arabinose. The growth was observed after incubation at 37°C for 24 h.

Subsequently, we analyzed the growth of strains expressing truncated *nbdA* variants. Strains producing NbdA variants that retained the MHYT domain showed similar behavior and failed to grow on agar plates upon induction. In contrast, production of truncated NbdA variants lacking the MHYT domain had no detectable effect on growth. To exclude the possibility that the observed growth defect resulted from general alteration of the cell membrane caused by overproduction of a membrane protein, three additional membrane proteins, MucR, BifA and CoxH were selected for control experiments. MucR and CoxH like NbdA, contain a triple MHYT domain at the N-terminus. Whereas MucR additionally harbors a cytosolic tandem DGC and PDE domain, CoxH contains a LytR domain (38). The overproduction of MucR, BifA and CoxH in PAO1 Δ*nbdA* did not affect growth on solid medium, regardless of the presence or absence of inducer (Fig. 3). These results indicate that the observed growth impairment is specific to NbdA production during the interaction with solid surfaces. Furthermore, we observed an elongation of cells in strains overproducing the MHYT-containing NbdA variants (Fig. S4).

To test whether this observed growth phenotype also results in impaired cell attachment, quantitative biomass attachment assays were performed with the *nbdA* overexpressor strains. Therefore, 96-well polystyrene plates were inoculated with strains overproducing NbdA variants and adhered biomass was determined after 1 h incubation at 37°C (Fig. 4). About 80% reduction in attached biomass was observed in this assay for the full length *nbdA* overexpressing strain compared to the strain harboring the empty vector (Fig. 4). Thereby, *nbdA* overexpression seemed to promote the planktonic lifestyle. For strains expressing truncated *nbdA* variants, we observed differences in the biomass adhesion depending on the truncation of protein domains (Fig. 3). The presence of the EAL domain, whether functional (EAL) or non-functional (AAL), resulted in a strong decrease in adhered biomass. The over-production of the MHYT domain alone, or together with the cytosolic AGDEF domain, increased biomass adhesion (Fig. 4).

**FIG. 4.**
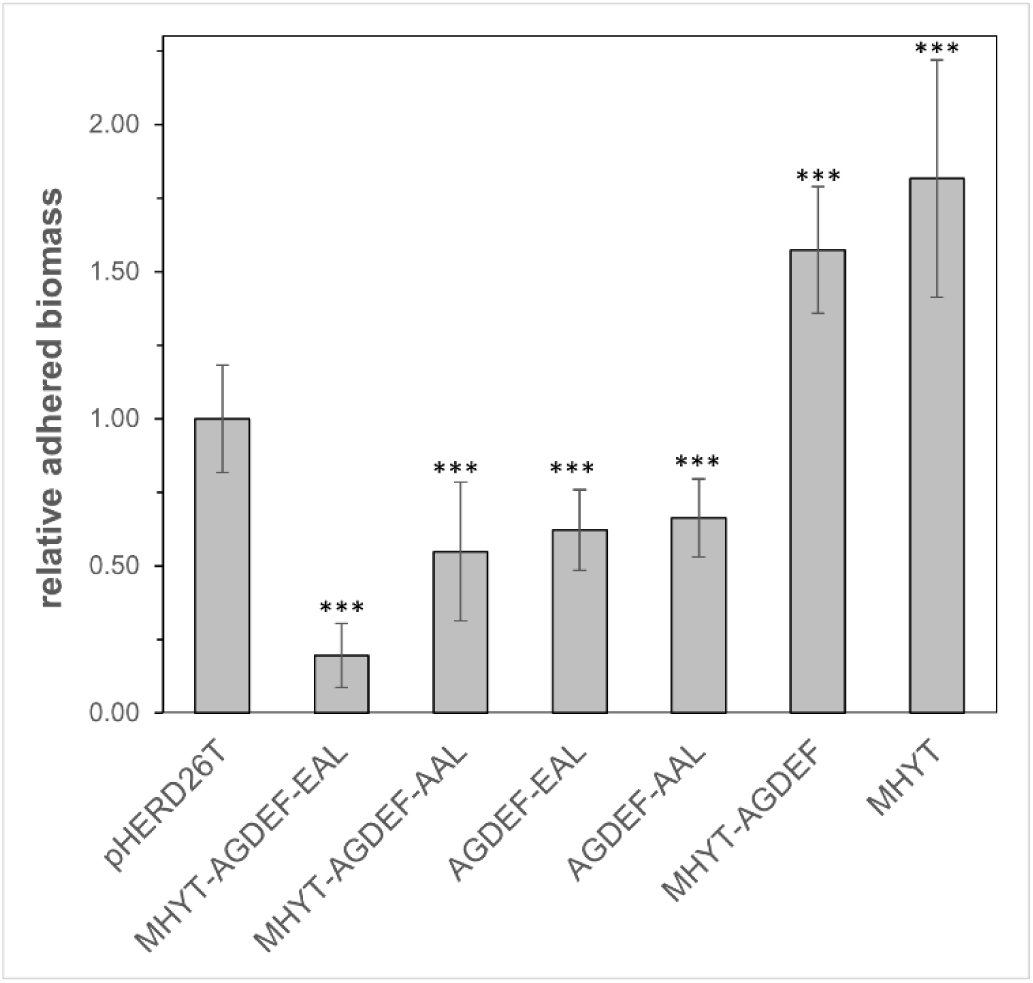
Quantitative attachment assay of *P. aeruginosa* and *nbdA* overexpressor strains. *P. aeruginosa* PAO1 *ΔnbdA* strains containing pHERD26T-derivatives for the overproduction of NbdA protein variants were grown in liquid BM2 with antibiotics. Gene expression was induced by adding 0.1% L-arabinose and further incubating for 5 h at 22°C. Adhesion to solid surfaces was tested in a polystyrene 96-well plate. For inoculation of the plate, cultures were adjusted to OD_600_ = 1.5 in adhesion medium. Each well was filled with 100 µL culture, and the 96-well plate was incubated for 1 h at 37°C in a moist chamber. Adhered biomass was determined by crystal violet staining. Adhesion was analyzed for each strain in three biological replicates with three technical replicates. Error bars represent the standard deviation. Significant changes are marked with asterisks, determined by the student’s t-test (*** p<0.001).

### Surface-dependent motility and type IV pili are affected by NbdA overproduction

The behaviour of NbdA overexpressor strains on solid surfaces led us to ask whether the formation or function of type IV pili (T4P) might be affected. We first analyzed twitching motility for the PAO1 *ΔnbdA* strain overexpressing *nbdA* and variants. The twitching motility was reduced strongly in the strain overexpressing full-length *nbdA* (Fig. 5). Interestingly, the PAO1 Δ*nbdA* strain producing the enzymatically inactive full-length variant of NbdA (NbdA-MHYT-AGDEF_AAL) was also unable to twitch, indicating a PDE-activity independent effect. To test whether overproduction of a membrane protein was responsible for the observed motility defect, twitching activity was determined for two control proteins BifA and CoxH mentioned earlier. CoxH overproduction in PAO1 did affect twitching motility only slightly, while the production of the phosphodiesterase BifA in PAO1 Δ*nbdA* reduced the twitching motility due to its PDE activity. These observations point towards a role of NbdA in surface-dependent motility, but independent of its PDE activity. Notably, strains overproducing truncated NbdA variants harboring the MHYT domain consistently displayed reduced twitching motility, while cytosolic variants without MHYT domain showed no significant impairment of twitching motility, regardless of whether the enzymatic domains were intact or catalytically inactive.

**FIG. 5.**
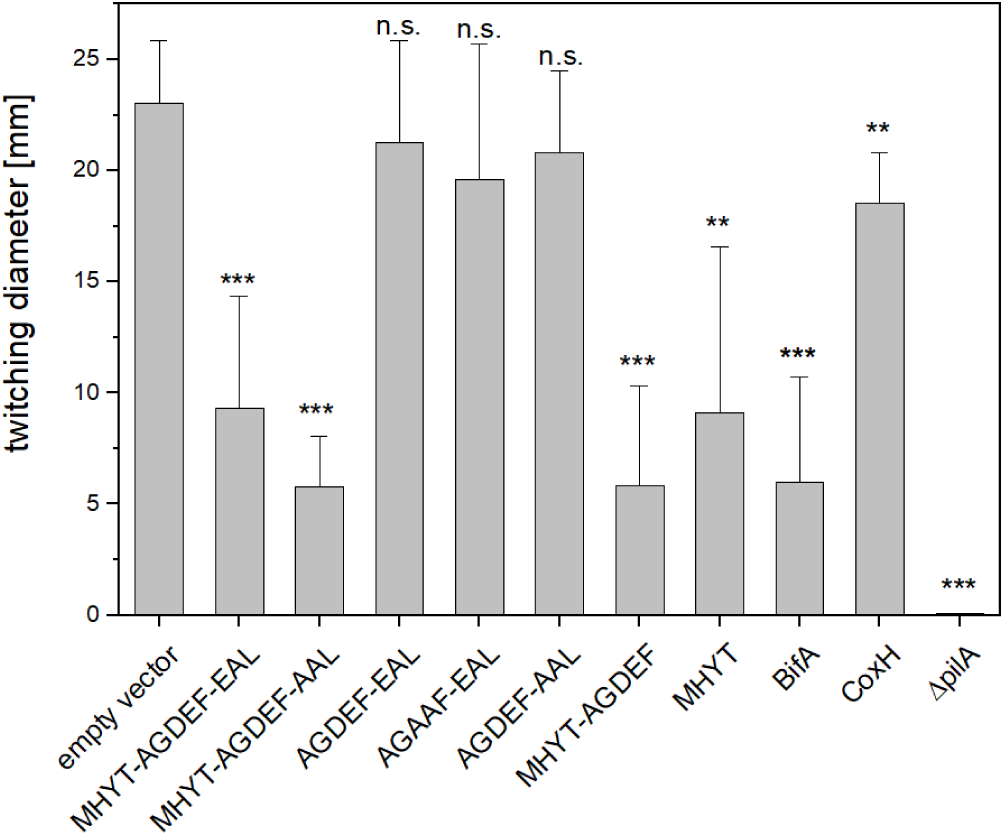
Twitching motility of PAO1 *ΔnbdA* complemented with plasmid-encoded NbdA fulllength and truncated variants. Cells were grown in LB and gene expression was induced at OD_600_ at 0.5 with 0.1% L-arabinose and incubated for 5 h at 37°C. Twitching plates were inoculated with 1 µL of culture (OD_600_ at 1.5) and incubated at 37°C and 42 h. Twitching zones were stained with 0.1% crystal violet solution. The experiment was performed at a minimum of three biological replicates with >4 technical replicates for each tested strain. Error bars represent the standard deviation. Significant changes are marked with asterisks, determined by the student’s t-test (*** p<0.001, ** 0.001 < *p* < 0.01, n.s. not significant).

### Overproduction of NbdA affects type IV pili-dependent phage infection

Twitching motility in *P. aeruginosa* is dependent on the formation and function of type IV pili (T4P). In order to test whether the observed phenotypes, namely the inability to grow on solid surfaces when *nbdA* is overexpressed, is attributable to a loss of T4P, a phage infection assay was employed. *P. aeruginosa* can only be infected by phage DMS3*vir* in the presence of functional T4P (36). PAO1 wildtype and PAO1 Δ*nbdA* strain showed an identical infection cycle and cell lysis when infected with DMS3*vir*, while the control strain PAO1 Δ*pilA* was not lysed by this phage. (Fig. S5).

Interestingly, PAO1 overexpressor strains showed normal lysis with DSM3vir in absence of inductor. When *nbdA* overexpression was induced in PAO1 prior to phage infection, the lysis curve was altered compared to the control strain (empty vector) and the uninduced culture (Fig. 6). Lysis of cells was significantly slower and less efficient than with the control strains. Furthermore, the number of released phage particles differed significantly. The PAO1 Δ*nbdA* harboring the empty vector released significantly more infective phage particles (2.7×10^12^ PFU/mL ± 2.1%) than PAO1 Δ*nbdA* strain overproducing NbdA (6×10^10^ PFU/mL ± 1.2%). These results suggest that overproduction of NbdA inhibited phage propagation, likely due to reduced T4 piliation or function.

**FIG. 6.**
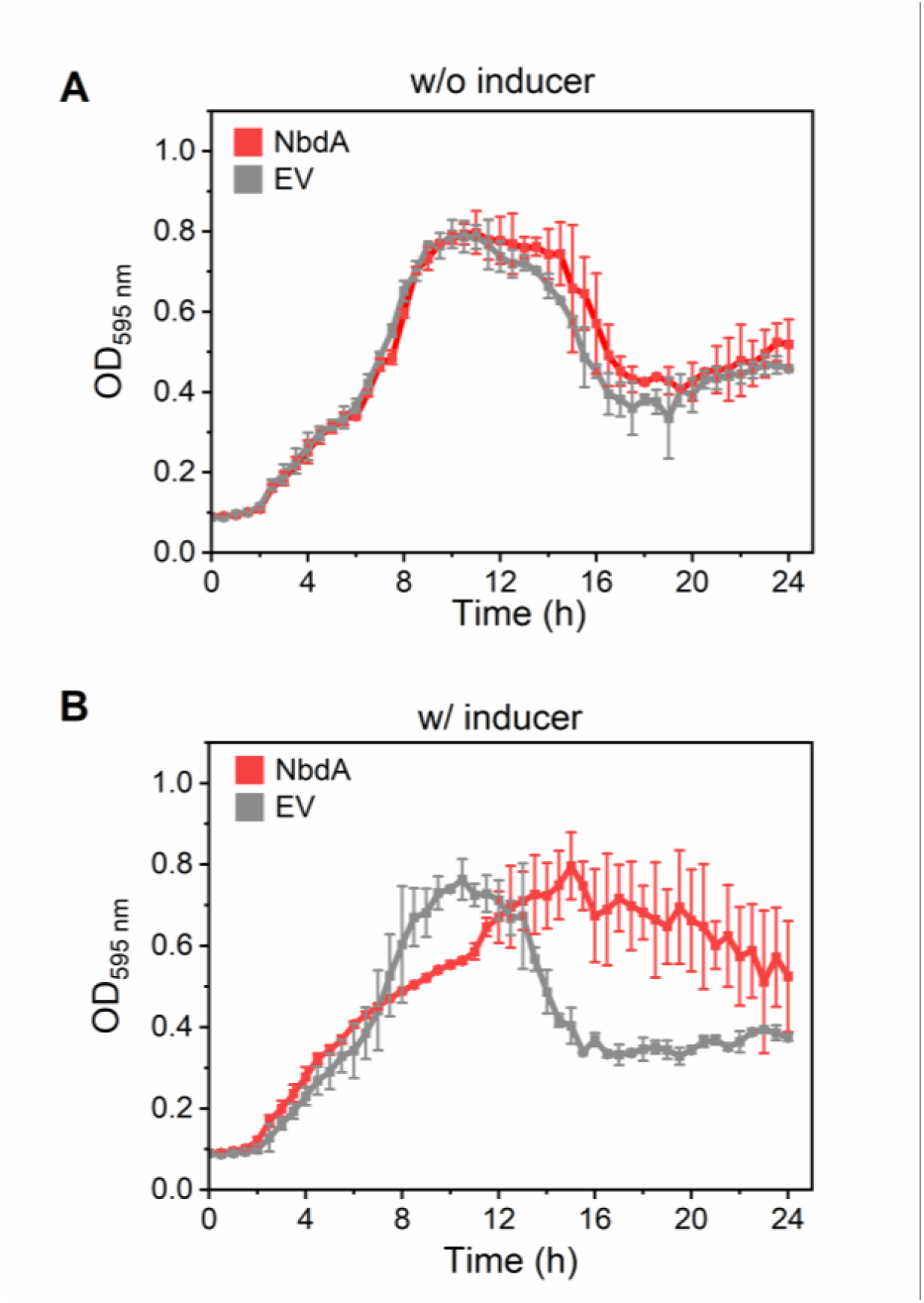
Phage infection assay with the phage DMS3vir on PAO1 Δ*nbdA* harboring the empty vector pME6032 (EV, grey) or the plasmid for overproduction of NbdA (NbdA, red). **A** Infection assay without induction of *nbdA* expression. **B** Infection assay with induction of *nbdA* expression. Cells were grown in a 96-well plate in LB media at 37°C. The OD_595_ was measured every 30 min in a plate reader (TECAN, Männedorf, Switzerland). Infection assays using 10^5^ PFU were performed in biological triplicates with three technical replicates each. Dots show the average of the biological triplicates; error bars indicate the standard deviation.

### NbdA has a dynamic localization in cells

Several of our results indicated an effect of NbdA on T4P formation or function. The localization of NbdA is required at the cell pole to enable interaction with T4P. We therefore studied the localization of NbdA using fluorescence microscopy. To this end, several fusion constructs of NbdA with the fluorescent proteins Venus and mNeonGreen were generated either as inducible and plasmid-encoded variants or genomically integrated in PAO1 at the original gene locus driven by its native promoter. Confocal microscopy images were acquired for both types of constructs. The cells with genomically integrated fusions showed a diverse fluorescence pattern. Lateral, polar and bipolar fluorescent foci were observed (Fig. 7). Furthermore, cells showed a diffuse fluorescence signal (38% and 50%) for NbdA-Venus and NbdA-mNeonGreen, respectively (Fig. 7). This was unexpected, as NbdA contains the MHYT membrane domain and would therefore be expected in the cell membrane. For the strains with the genomic integration of *nbdA-*fluorescent gene fusions we were unable to prove the protein production by western blot. Thus, these strains were not further used. For cells with plasmid-encoded and induced NbdA-fusions, we observed fluorescent foci at one cell pole (43%) and foci at both cell poles (6%) (Fig. 7). Still, about 50% of the cells showed a diffuse fluorescent signal. This indicated that NbdA was dynamically distributed in the cell with partial accumulation at the cell pole.

**FIG. 7.**
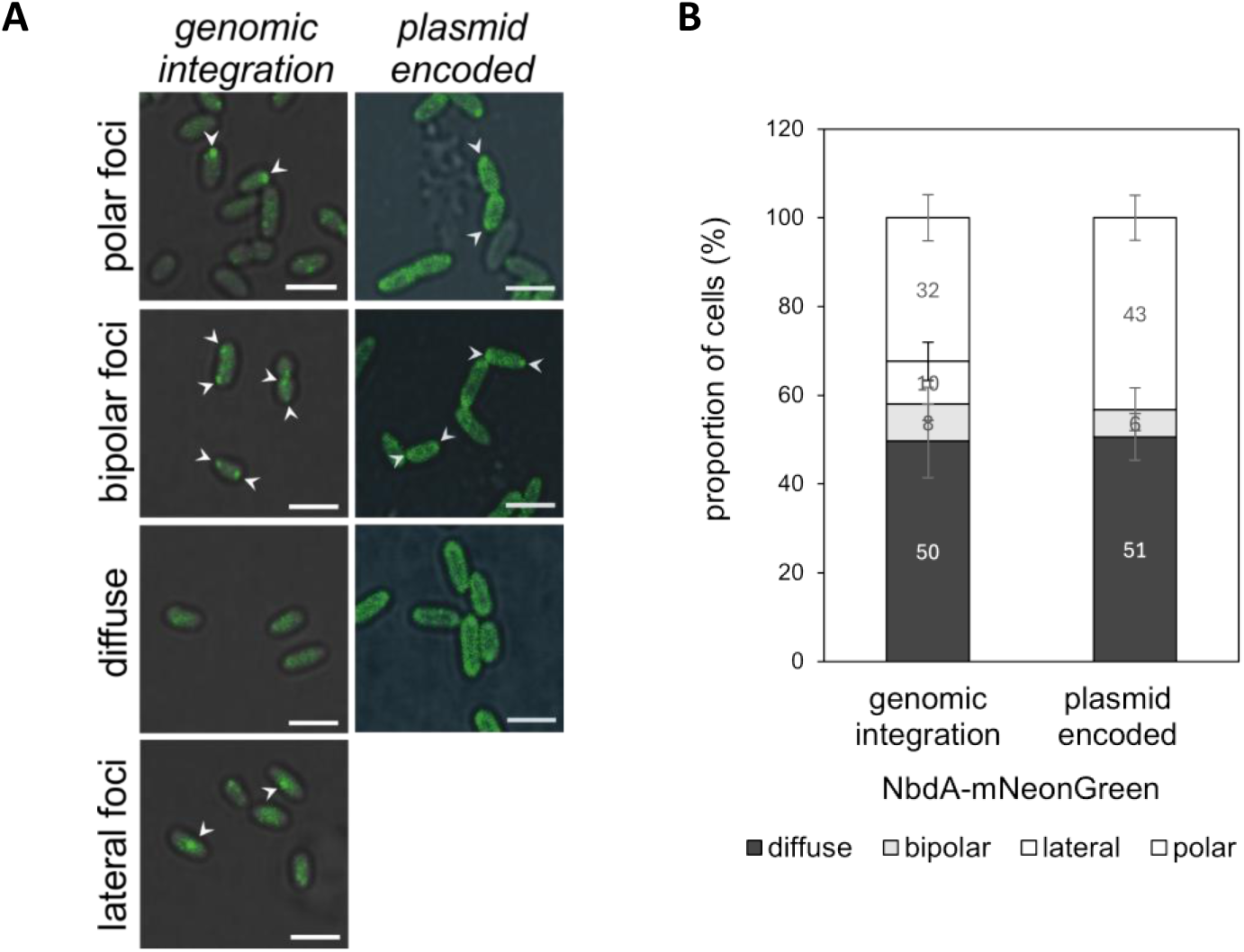
Localization of NbdA-mNeonGreen in PAO1. **A**: Micrographs of PAO1 carrying a plasmid encoded copy of NbdA-mNeonGreen (merged differential interference contrast and fluorescence image). Examples for the observed signal patterns are shown. Arrowheads indicate the positions of fluorescent foci for NbdA-mNeonGreen. Scale bar: 2 μm. **B**: Proportion of NbdA-mNeonGreen signals in the observed cells are depicted. Expression of NbdA was controlled either from the native promoter or from the inducible tac promoter (pME6032). Error bars indicate the standard deviation from three independent experiments. Total cell counts: NbdA (native promoter): *n* = 1101, NbdA (pME6032): *n* = 1395.

The cell pole harbors essential motility related structures like flagella and T4P. Previous studies have linked NbdA to T4P, the flagellar system, and the chemotaxis machinery (24–26). Therefore, we investigated the co-localization of NbdA with components of the T4P system, the flagellar system, and the chemotaxis system using fluorescence microscopy. The subunit FliC (T394C) of the flagellum was labelled with Alexa-fluor 594 maleimide and was detected exclusively at one cell pole. Co-localization experiments with NbdA-mNeonGreen showed different localization patterns (Fig. 8). About 47% of cells showed fluorescent foci for NbdA-mNeonGreen at the same cell pole as the flagella. However, one third of the observed cells (32%) showed a diffuse distribution of fluorescence for NbdA-mNeonGreen on the inner membrane. Furthermore, but less prominently, some NbdA-mNeonGreen fluorescence was distributed at the opposite cell pole to the flagella FliC-signal, and at both cell poles simultaneously (Figure 8). In summary, nearly half of the cells showed NbdA-mNeonGreen localized to the flagellated cell pole.

**FIG. 8.**
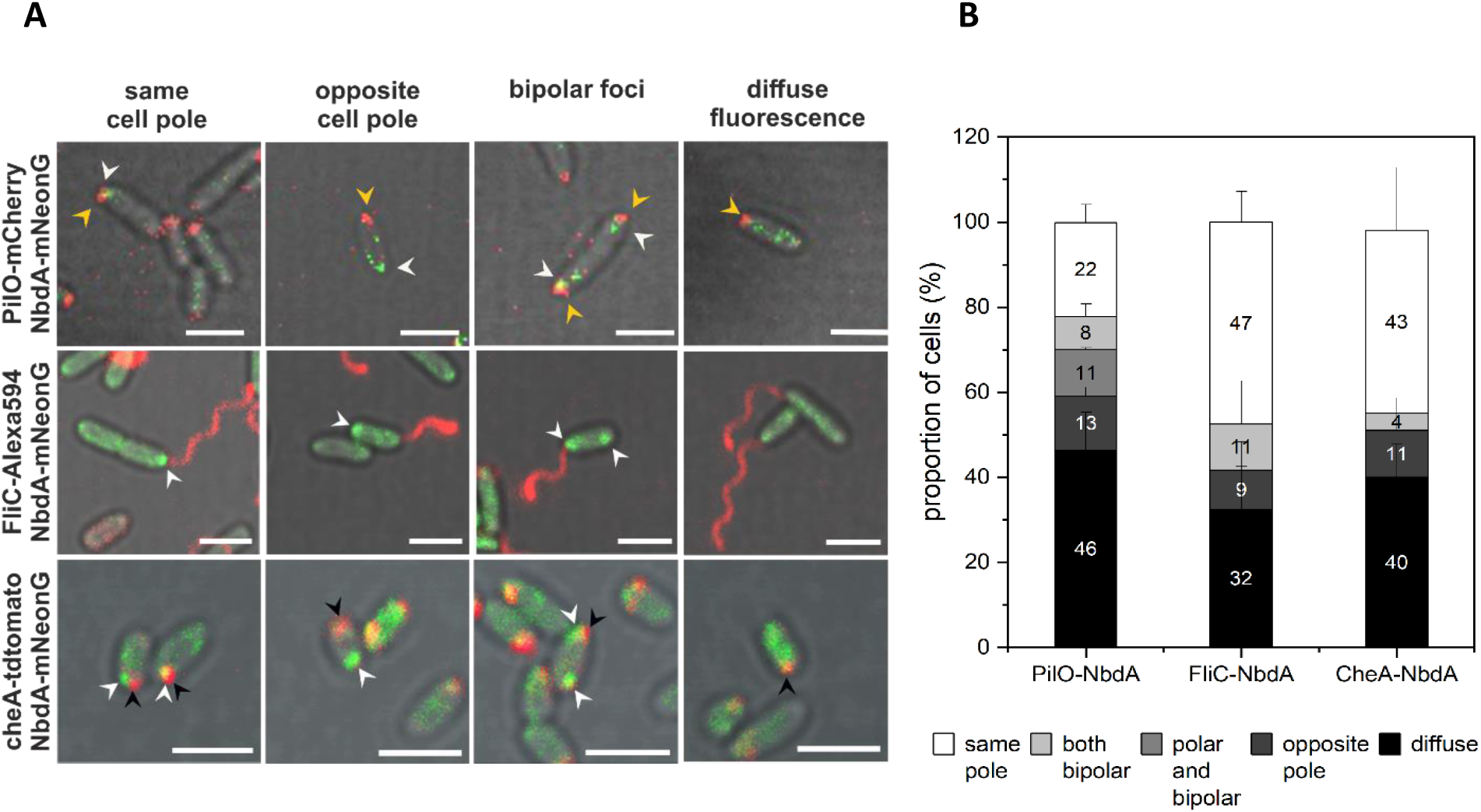
Co-localization tests of NbdA with the type IV pilus, flagella, and the chemotaxis apparatus. **A**: Micrographs showing NbdA-mNeonGreen patterns (green) observed in combination with type IV pilus component PilO-mCherry (red, top panel), with Alexa594 stained flagella subunit FliC (red, middle panel) and the chemotaxis protein CheA-tdTomato (red, lower panel). Merged images from differential interference contrast and fluorescence are shown. Flagella were stained by labelling FliC_T394C with the fluorescent dye Alexa Fluor™ 594 C5 Maleimide (red). NbdA-mNeonGreen foci are indicated with white arrow heads, CheA-tdTomato with black arrow heads, and PilO-mCherry with yellow arrow heads. Scale bar: 2 μm. **B**: Distribution of fluorescence signals in the cells for localization of NbdA with CheA, FliC or PilO. Depicted are the proportions of cells for each signal combination that was observed. Error bars indicate the standard deviation. Total cell counts: CheA and NbdA: *n* = 856, FliC and NbdA: *n* = 293, PilO and NbdA: *n* = 166.

Detection of the T4P apparatus was achieved by a PilO-mCherry fusion. PilO is a structural compound of the intact T4P at the inner membrane (39). PilO-mCherry showed fluorescence in polar, and less frequently, in bipolar foci. The distribution of fluorescent foci for NbdA-mNeonGreen and PilO-mCherry in cells was classified into three groups (Fig. 8). In Group I, representing 30% of the cells, PilO-mCherry and NbdA-mNeonGreen were co-localized in polar foci (22%) and bipolar foci (8%). Group II, with 15% of the cells, polar foci for PilO-mCherry and bipolar foci for NbdA-mNeonGreen were detected in the cells. Group III, consisted of cells showing distinct foci but no co-localization. In this group, foci for NbdA and PilO protein fusions were detected at opposite poles (13%). Further 42% of cells showed diffuse fluorescence distribution for NbdA-mNeonGreen. Therefore, these cells were not assigned to any of the previous groups. These observations showed the dynamic nature of NbdA distribution in the cell. When NbdA was found in the cell membrane, the T4P protein PilO co-localized in the majority of cells.

Co-localization of NbdA with the chemotaxis system was tested with the fusion construct of CheA-tdTomato. Around 43% of cells show fluorescence signals for NbdA-mNeonGreen and CheA-tdTomato at the same cell pole. A similar number of cells (40%) show diffuse fluorescence for NbdA-mNeonGreen. The remaining cells (17%) show disparate localization patterns, either lateral or bipolar for NbdA; or distribution at the opposite cell pole compared to CheA-tdTomato.

## Discussion

In this work, we demonstrated the PDE activity of the full-length protein NbdA *in vivo* through heterologous complementation of the *pdeH* deletion in *E. coli* AB607. This extends the previous results, the *in vitro* activity of the protein’s cytosolic domains (22). Furthermore, it confirmed that the PDE domain of NbdA is responsible for phosphodiesterase activity (22). In contrast to former *in vitro* experiments, the truncated NbdA variant of the standalone EAL domain showed no sign of *in vivo* PDE activity. Complementation of the *E. coli* ΔpdeH phenotype occurred only with the combined DGC and PDE domains. This observation suggested an interference or regulatory role of the DGC domain over the PDE activity in NbdA. Several dual-domain proteins have been described where one domain controls the activity of the other. An allosteric control was described for the tandem DGC-PDE protein CC3396 of *Caulobacter crescentus*, where GTP binds to the DGC-domain and thereby promotes the activity of the PDE domain (40). The *in vitro* PDE activity of cytosolic NbdA was also shown to be enhanced by GTP (22). Further, protein conformation and rearrangement, like in the membrane bound DGC-PDE tandem protein RbdA from *P. aeruginosa*, can affect substrate binding (41, 42). But not only intramolecular regulation is crucial for activity *in vivo*. The binding of signal molecules and the interaction with local target proteins are the most prominent arguments for the control of enzymatic activity *in vivo* (20, 21, 43–51). To confirm the enzymatic function of NbdA in its native host, we determined the cellular concentration of the nucleotides c-di-GMP, pGpG, and GMP by HPLC-ESI_pos_-MS/MS using sMRM simultaneously in cell extracts of PAO1 wildtype, PAO1 Δ*nbdA*, and PAO1 overexpressing plasmid-encoded variants of *nbdA*. The c-di-GMP content of cell extracts from wildtype and the *nbdA* deletion strain (PAO1 Δ*nbdA*) was similar and consistent with previous studies (23, 24, 52). This implies a rather local role of NbdA than a global function in cellular c-di-GMP degradation. The overexpression of full-length *nbdA* resulted in slight but not strongly reduced c-di-GMP levels. In contrast, the production of truncated cytosolic and modified protein variants of NbdA in *Pseudomonas* showed increased levels of c-di-GMP. This was a surprising result, as we did not expect DGC activity for NbdA. The protein sequence harbors a degenerated DGC motif (AGDEF) that was considered to be enzymatically inactive and no DGC activity of NbdA has been observed previously (22). Although, the canonical “GGDEF” motif was shown to be essential, situated in the active site of DGC proteins (8, 40, 53), some diguanylate cyclase proteins, e.g. VCA0965 of *Vibrio cholerae* and XAC0610 of *Xanthomonas citrici* contain the degenerate “AGDEF” motif and are active enzymes (54, 55). Changing the canonical GGDEF motif to AGAAF or GEDEF in other DGC proteins made the respective enzymes inactive (40, 56). Still, *Pseudomonas* cells producing the cytosolic NbdA-AGAAF-EAL variant showed increased c-di-GMP levels. The overproduction of the cytosolic domains and not an enzymatic activity is likely responsible for this effect since overproduction of the full-length protein or the membrane MHYT domain alone did not increase the c-di-GMP levels. Protein-interactions, heterodimerization, or activation of other c-di-GMP modulating enzymes are possible reasons for the observed disbalance of c-di-GMP level in the cells. Large modules of DGCs, PDEs and effector proteins were shown to control local cellular functions, while only few PDEs and DGCs are responsible for the overall cellular c-di-GMP homeostasis (12, 13, 17, 19, 21, 57–59). In several hosts, strong heterodimer formation between DGCs and PDEs have been observed that result in modulation of their enzymatic activity (12, 19). The diguanylate cyclase-phosphodiesterase pair YciR (PdeR) and YdaM (DgcM) in *E. coli* is regulating enzymatic activities by interaction (57). Perception of different signals can also regulate enzyme activity. For example, the DGC activity of SadC is inhibited by interaction with PilO of the pilus alignment complex PilMONP, and MotC of the flagellar motor (13, 21, 60). Thereby it integrates surface signals derived from the flagellar system and T4P to coordinate surface attachment (21, 59). SadC was also previously found to interact with NbdA (19, 26). Furthermore, NbdA has been linked to various processes involved in biofilm formation and motility (22, 24, 29). In *P. aeruginosa* KT1115 the homolog protein was found to coordinate exopolysaccharid production and rhamnolipid synthesis (29). In PAO1, NbdA was shown to be involved in chemotaxis-mediated flagellar motor switching (24). Furthermore, direct protein interaction of NbdA with the T4P extension motor PilB and several T4P components (PilD, PilU, PilO, PilY2) was confirmed (26). Similarly to T4P modulating proteins SadC and DgcP, NbdA might be part of the network to modulate c-di-GMP levels involved in motility and T4P mediated surface behaviour (7, 10, 13, 14, 21, 61, 62). The deletion strain PAO1 Δ*nbdA* did not exhibit a strong biofilm or motility phenotype, but the overproduction of NbdA resulted in growth inhibition on solid surfaces and reduced surface attachment and twitching activity, which suggests interference with T4P. This hypothesis was further supported by the infection assay with the T4P-dependent phage DMS3vir. The strain overproducing NbdA was no longer successfully infected by this phage. This indicates a defect in T4P production, or function. Our study revealed a dynamic distribution of NbdA in cells which is similar to the cellular distribution of fluorescent signals that was previously observed for GFP-PilT, the pilus retraction motor (39, 63). The pilus extension and retraction motors PilB, and PilT, respectively, bind competitively to pilus machineries, and could be used as a reporter for active T4P machines in *P. aeruginosa* (39). PilB is not always bound to the pilus machinery, therefore we used PilO-mCherry to detect T4P co-localization patterns with NbdA. The interaction of NbdA with PilB and PilO was previously shown in pull down analysis (26). Fully assembled pilus machines were shown at both cell poles (39), yet a strict colocalization of both proteins, PilO and NbdA could not be confirmed in this study. In a cell, several active and inactive T4P machines are present. The pilus extension and retraction are highly dynamic and are activated depending on different input signals like cAMP and c-di-GMP (39). We propose a role for NbdA in the regulation network of surface dependent behavior and T4P function by protein-protein interactions and modulation of c-di-GMP in *Pseudomonas aeruginosa*.

## Acknowledgements

We thank Miriam Haak for technical assistance. This work was supported within the German research foundation framework of the priority program SPP1879 to NFD.

